# Recombination in Sarbecovirus lineage and mutations/insertions in spike protein linked to the emergence and adaptation of SARS-CoV-2

**DOI:** 10.1101/2020.05.12.091199

**Authors:** Amresh Kumar Sharma, Priyanka Kumari, Anup Som

## Abstract

The outbreak of severe acute respiratory syndrome coronavirus 2 (SARS-CoV-2) in Wuhan city, China in December 2019 and thereafter its spillover across the world has created a global pandemic and public health crisis. Researchers across the world are involved in finding the origin and evolution of SARS-CoV-2, its transmission route, molecular mechanism of interaction between SARS-CoV-2 and host cells, and the cause of pathogenicity etc. In this paper, we shed light on the origin, evolution and adaptation of SARS-CoV-2 into human systems. Our phylogenetic/evolutionary analysis supported that bat-CoV-RaTG13 is the closest relative of human SARS-CoV-2, outbreak of SARS-CoV-2 took place via inter-intra species mode of transmission, and host-specific adaptation occurred in SARS-CoV-2. Furthermore, genome recombination analysis found that Sarbecoviruses, the subgenus containing SARS-CoV and SARS-CoV-2, undergo frequent recombination. Multiple sequence alignment (MSA) of spike proteins revealed the insertion of four amino acid residues “PRRA” (Proline-Arginine-Arginine-Alanine) into the SARS-CoV-2 human strains. Structural modeling of spike protein of bat-CoV-RaTG13 also shows a high number of mutations at one of the receptor binding domains (RBD). Overall, this study finds that the probable origin of SARS-CoV-2 is the results of intra-species recombination events between bat coronaviruses belonging to Sarbecovirus subgenus and the insertion of amino acid residues “PRRA” and mutations in the RBD in spike protein are probably responsible for the adaptation of SARS-CoV-2 into human systems. Thus, our findings add strength to the existing knowledge on the origin and adaptation of SARS-CoV-2, and can be useful for understanding the molecular mechanisms of interaction between SARS-CoV-2 and host cells which is crucial for vaccine design and predicting future pandemics.

## 1. Introduction

Coronaviruses are single-stranded RNA viruses of 26 to 32 kilobases (kb) nucleotide chain and consist of both structural and non-structural proteins. They have been known to cause lower and upper respiratory diseases, central nervous system infection and gastroenteritis in a number of avian and mammalian hosts including humans (Zhu et al., 2019; Gorbalenya et al 2020). The recent outbreak of novel coronavirus (SARS-CoV-2) associated with acute respiratory disease called coronavirus disease 19 (commonly known as COVID-19) has caused a global pandemic. As of 15^th^ June 2021, more than 175 million laboratory confirmed COVID-19 cases and approximately 3.78 million people have died and further COVID-19 appears as a global threat to public health as well as to the human civilization as economic and social disruption caused by the pandemic is devastating (WHO, COVID-19 situation reports).

Coronaviruses are placed within the family *Coronaviridae*, which has two subfamilies namely *Orthocoronavirinae* and *Torovirinae. Orthocoronavirinae* has four genera: *Alphacoronavirus* (average genome size 28kb), *Betacoronavirus* (average genome size 30kb), *Gammacoronavirus* (average genome size 28kb), and *Deltacoronavirus* (average genome size 26kb) (King et al. 2011). Coronaviruses are typically harbored in mammals and birds. Particularly *Alphacoronavirus* and *Betacoronavirus* infect mammals, and *Gammacoronavirus* and *Deltacoronavirus* infect avian species (Woo et al., 2009; 2010; Fan et al., 2019). SARS-CoV-2 is a member of the genus *Betacoronavirus* and subgenus Sarbecovirus. Figure 1 shows the taxonomical origin of SARS-CoV-2.

**Figure 1:**
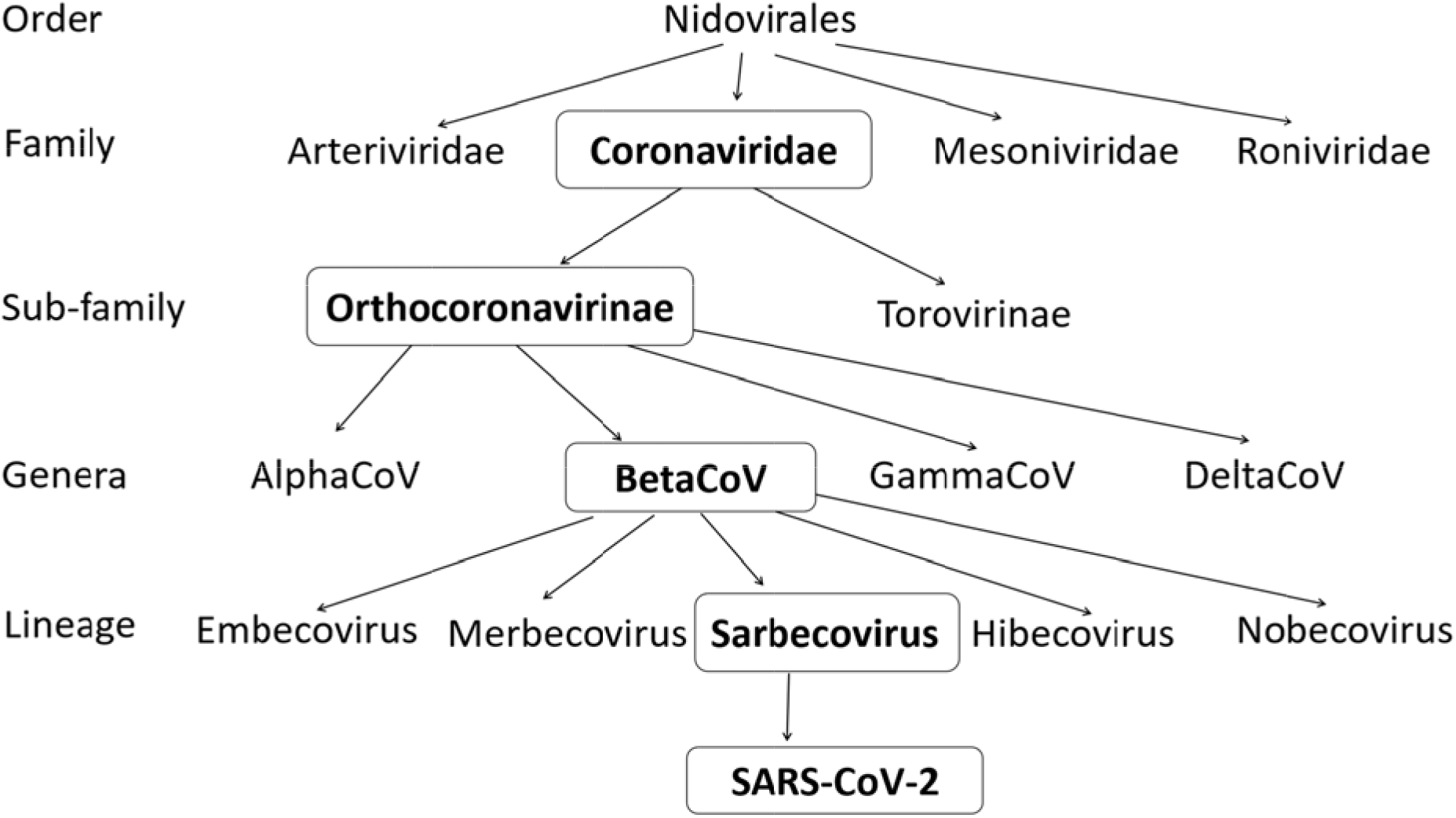
Taxonomical origin/classification of SARS-CoV-2.

The previous important outbreaks of coronaviruses are severe acute respiratory syndrome coronavirus (SARS-CoV or SARS-CoV-1) outbreak in China in 2002/03, Middle East respiratory syndrome coronavirus (MERS-CoV) outbreak in 2012 that resulted severe epidemics in the respective geographical regions (Eickmann et al., 2003; Vijaykrishna et al., 2007; Zumla et al, 2015; Hayes et al., 2019). The present outbreak of SARS-CoV-2 is the third documented spillover of an animal coronavirus to humans in only two decades that has resulted in a major pandemic (Velavan and Meyer, 2020; Lai et al., 2020; Srivastava et al., 2021).

Scientific communities across the world are trying to understand several fundamental and applied questions such as: What is the origin of SARS-CoV-2? How does SARS-CoV-2 adapted to infect humans? What are the possible transmission routes? Why SARS-CoV-2 is more deadly than other CoVs? What is its possible clinical diagnosis & treatment? etc. Consequently, a large number of research outcomes are being consistently published. In this paper, we aim to shed light on the origin, evolution and adaptation of SARS-COV-2 using molecular phylogeny, evolutionary and structural modeling studies.

## 2. Materials and Methods

### 2.1. Data selection

162 *Orthocoronavirinae* genomes were retrieved from NCBI (https://www.ncbi.nlm.nih.gov/) and Virus Pathogen Database and Analysis Resource (https://www.viprbrc.org/). We only considered complete genome sequences having no unidentified nucleotide characters. Our dataset included 23 *Alphacoronavirus*, 92 *Betacoronavirus*, 32 *Deltacoronavirus* and 15 *Gammacoronavirus* genomes belonging to different subgenus, diverse host species and from wide geographical location. Further for rooting the tree, we used two genome sequences from *Torovirus* and two from *Bafinivirus* belonging to domestic cow and fish respectively. The genera *Torovirus* and *Bafinivirus* belong to the sub-family *Torovirinave* of the family *Coronaviridae.* Overall, the phylogenetic analysis consists of 166 complete viral genomes (162 *Orthocoronavirinae* and four *Torovirinave genomes*). Details genome sequences used in this study can be found in Supplementary File S1.

### 2.2. Phylogenetic reconstruction

The genome sequences were aligned using the MAFFT alignment tool (Katoh et al., 2002). Genome tree of the *Orthocoronavirinae* and *Betacoronaviruses* were reconstructed using maximum likelihood (ML) method and GTR+G+I model of nucleotide substitution as revealed by the model test with 1000 bootstrap support. The model test was performed for accurate phylogenetic estimation by using ModelFinder, which is implemented in IQ-TREE version 1.5.4 (Kalyaanamoorthy et al., 2017). Phylogenetic trees were reconstructed using IQ-TREE software (Nguyen et al., 2015). The trees were visualized with iTOL tool (Letunic et al., 2019). Five gene trees of the *Betacoronaviruses* were reconstructed using Orf1ab, Spike (S), Envelope (E) Membrane (M), and Nucleocapsid (N) amino acid sequences. The ML method of tree reconstruction and protein-specific amino acids substitution model as revealed by ModelFinder was used for gene tree reconstruction. Bootstrap test with 1000 bootstrap replicates was carried out to check the reliability of the gene trees.

### 2.3. Genome and gene recombination analysis

Potential recombination events in the history of the *Betacoronaviruses* were assessed using the RDP5 package (Martin et al., 2015). The RDP5 analysis was conducted based on the complete genome sequence using RDP, GENECONV, BootScan, MaxChi, Chimera, SiScan, and 3Scan methods. Putative recombination events were identified with a Bonferroni corrected P-value cutoff of 0.05 supported by more than four methods.

### 2.4. Sequence and structural analysis

The homology and genetic variations analysis of sequences in different genomic regions of SARS-CoV-2 strain Wuhan Hu-01 (MN908947) is compared to bat-CoV-RaTG13 (MN996532) and pangolin-CoV-GX-P5E (MT040336) using CLUSTAL W (https://www.genome.jp/tools-bin/clustalw) and multiple sequence alignment (MSA) analysis of spike proteins were performed using CLUSTAL OMEGA (https://www.ebi.ac.uk/Tools/msa/clustalo/).

The structures of the spike protein of SARS-CoV-2 Wuhan Hu-1 (PDB: 6XLU), bat-CoV-RaTG13 (PDB: 6ZGF) were retrieved from PDB database (Rose et al. 2016). The spike protein for pangolin coronavirus was not available so it was modeled using SWISS-MODEL SERVER (https://swissmodel.expasy.org) with 6XR8 as template. These structures were compared using the structure superimposition/structure alignment tool of Chimera software (Pettersen et al. 2004).

## 3. Results and Discussion

In this study we aim to understand the origin and evolutionary trajectory of SARS-CoV-2 using molercular phylogenetic, genetic recombination and structural analyses. Particularly, we study the origin of SARS-CoV-2 from their deep ancestral roots (i.e., from deeper shared evolutionary history). Accordingly, the molecular phylogenetic analysis was based on two-stage genome phylogeny followed by gene trees analyses. Firstly, reconstruction of genome phylogeny of the *Orthocoronavirinae* genomes and study the cladistic/evolutionary relationships of its four genera. Secondly, reconstruction of *Betacoronavirus* genome and gene phylogeny that included its five subgenera namely Embecovirus, Hibecovirus, Merbecovirus, Nobecovirus and Sarbecovirus, and study the evolutionary relations of these five subgenera. The genome phylogeny of *Orthocoronavirinae* depicts that Alpha, Beta, Delta and Gamma coronaviruses clustered according to their cladistic relations where *Alphacoronavirus* clade appeared as a basal radiation of the *Orthocoronavirinae* phylogeny (Fig. 2). This result is consistent with the other results (Luk et al. 2019; Wu et al., 2020). Furthermore, analysis of the clades found that *Gammacoronavirus* and *Deltacoronavirus* clades are monophyletic (originated from a single common ancestor). This result is supported by their hosts’ nature; as both types mostly infect avian species (Wertheim et al. 2013).

**Figure 2:**
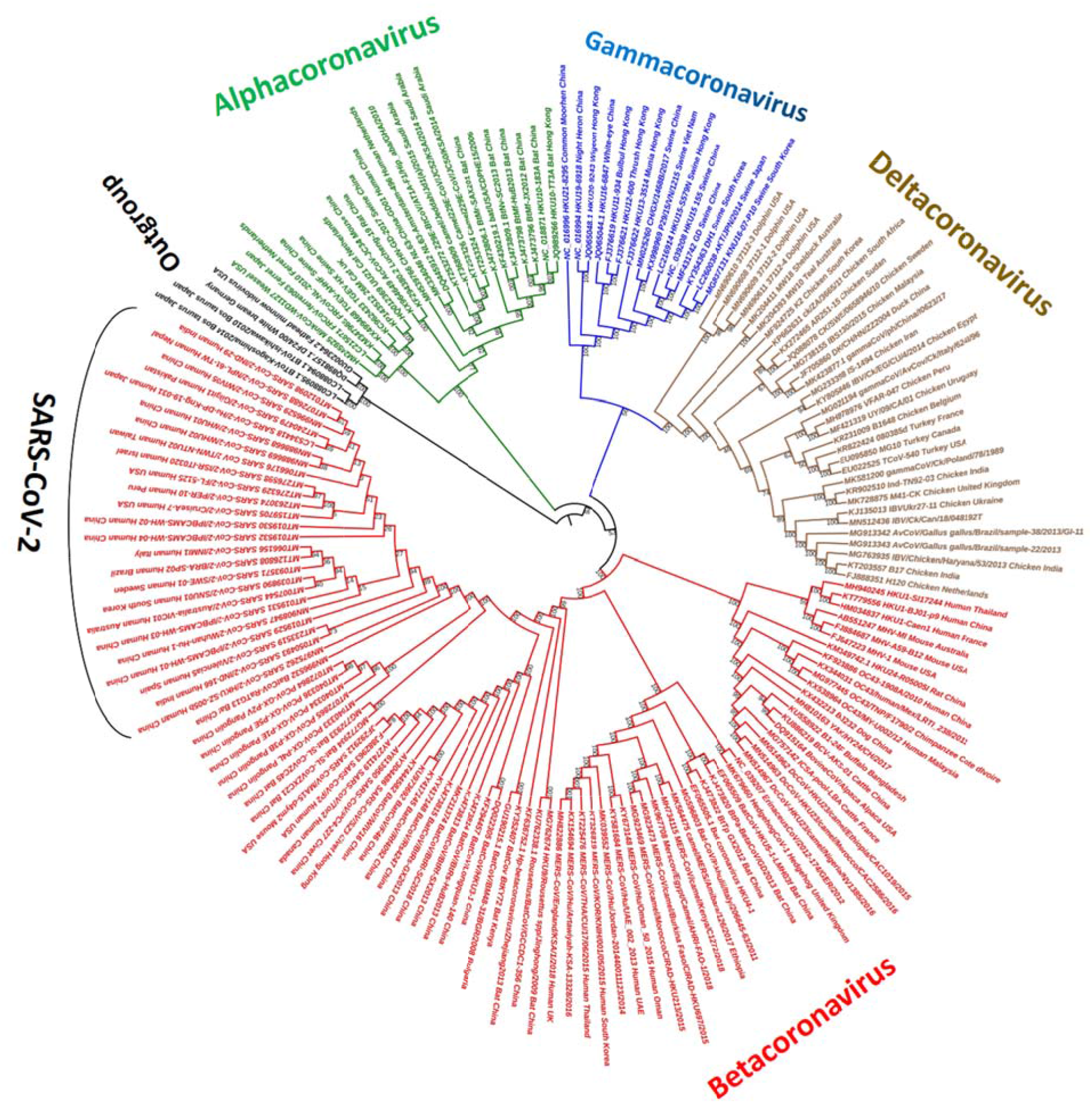
*Orthocoronavirinae* genome phylogeny. The genome tree consists of 162 complete *Orthocoronavirinae* genomes and four outgroups. Alignment consists of 58,538bp aligned nucleotide characters (9,384bp are completely aligned characters). Tree was reconstructed using ML method with GTR+G+I model of nucleotide evolution along with 1000 bootstrap replicates. Tree was rooted with the four Torovirinae genomes (outgroup). SARS-CoV-2 genomes are depicted in *Betacoronavirus*.

Further, a deeper analysis of the *Orthocoronavirinae* genome tree revealed that irrespective of their geographical locations, the host-specific strains are clustered together.This is probably due to the host adaptation, which is an important characteristic of viral genomes for their survival and replication (Songa et al., 2005; Fung et al., 2019; Andersen et al., 2020).

For example, *Alphacoronavirus* strains from ferret_Japan and ferret_Netherland are monophyletic. Similarly cat_UK is monophyletic with cat_Netherland, and human_China is monophyletic with human_Netherland. Further analysis revealed all *Alphacoronavirus* camel strains of Saudi Arabia appeared in a distinct subclade where bat_Ghana strain appeared as outgroup which indicates interspecies transmission took place from bat_Ghana to camel. A body of literature also reported that SARS-CoV-2 transmission took place to humans through intermediate hosts (Montoya et al., 2020; Roy et al., 2021; York, 2020; Zhou et al., 2020).

Deltacoronavirus and Gammacoronavirus clades exhibit a similar evolutionary pattern. In case of Deltacoronaviruses, swine_Vietnam and swine_Hong Kong shared a single common ancestor. Similarly, swine_China and swine_South Korea are monophyletic clade and swine_Japan is monophyletic with swine_South Korea.

In case of *Gammacoronaviruses* (whose natural hosts are avian species), chicken Peru and chicken_Uruguay are monophyletic. Similarly, chicken_Iraq is monophyletic with chicken_Egypt strain. These results reconfirm that coronaviruses are present in a large number of hosts those are widespread in different geographical location and coronaviruses undergo host-specific adaptation (Nakagawa and Miyazawa, 2020).

Phylogenetic analysis of *Betacoronavirus* genomes revealed that the five subgenera clustered separately (Fig. 3). Furthermore, the *Betacoronavirus* genome tree depicts that the host-specific strains from distance geographical locations formed monophyletic clades. For example, in Embecovirus clade, strain BJ01_P9_human_China is monophyletic with Caenl_human_France strain. Similarly, Embecovirus B1_24F_buffalo_Bangladesh is monophyletic with BCV_AKS_01_cattle_China. This result suggests host adaptation of SARS-CoV-2 had occurred (Fung et al., 2019; Montoya et al., 2020; Roy et al., 2021).

**Figure 3:**
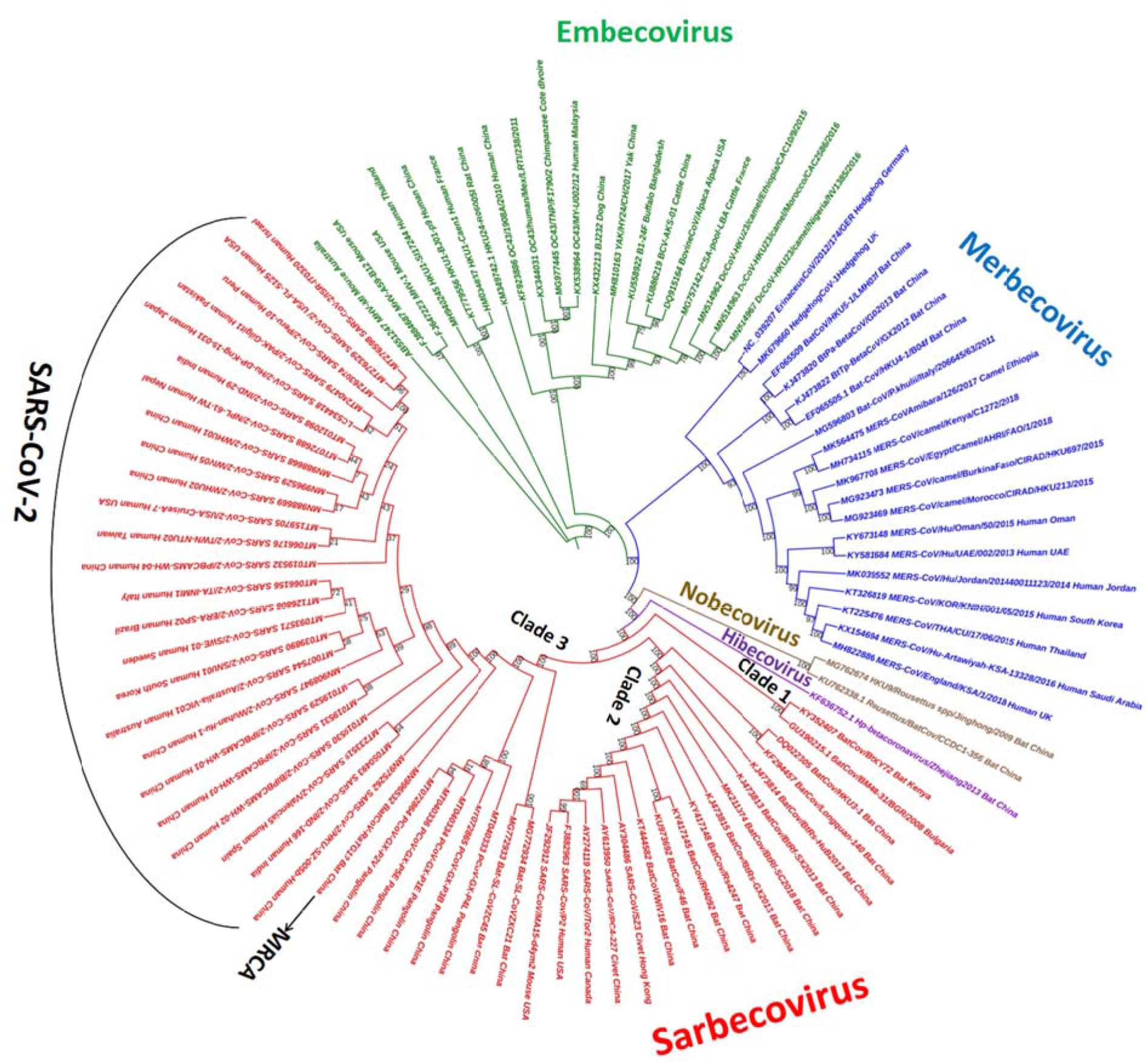
*Betacoronavirus* genome phylogeny. The genome tree consists of 92 complete *Betacoronavirus* genomes. Alignment consists of 41,054bp aligned nucleotide characters (23,064 bp are completely aligned characters). Tree was reconstructed using ML method with GTR+G+I model of nucleotide evolution along with 1000 bootstrap replicates. Most recent common ancestor (MRCA) of SARS-CoV-2 is highlighted. Three distinct clades of Sarbecovirus are also depicted.

SARS-CoV-2 belongs to Sarbecovirus subgenus. Sarbecoviruses formed three distinct clades (Fig. 3), where Clade 1 consists of only bat as host species. In Clade 2, host species are bat, civet and human. Similarly, in Clade 3 the host species are bat, pangolin and human and it depicts bat-CoV-RaTG13 (NCBI Acc no. MN996532) is closest to the human SARS-CoV-2 as all human SARS-CoV-2s clustered in a clade, and formed a monophyletic clade to bat-CoV-RaTG13 strain (i.e. descended from a common ancestor). Clade 3 also shown that pangolin (PCoV-GX-P5E) is the second closest relative of human SARS-CoV-2 behind bat-CoV-RaTG13. This result was also reported by other studies (Liu et al., 2020; Zhang et al., 2020). Further, deep node analysis, in Clade 3, suggested that SARS-CoV-2s, pangolin CoVs (strains PCoV-GX-P4L/P3B/P1E/P5E/P2V) and bat-CoVs (strains bat-SL-CoVZXC21 and bat-SL-CoVZC45) shared a single common ancestor (Fig. 3). These clades analysis suggest bat and pangolin are the natural reservoir of SARS-CoV-2 and possibly transmission from bat /pangolin to humans took place through intermediate organisms.

This similar observation had also been reported by a body of literature (Cui et al., 2019; Fung et al., 2019; Montoya et al., 2020; Roy et al., 2021; York, 2020). Furthermore, phylogenetic analysis reveals that the *Betacoronavirus* sequences including SARS-CoV-2s are conserved in their respective hosts (e.g. all bat hosts clustered in Clade 2 and human hosts are in Clade 3). This is probably due to host-specific adaptation to facilitate colonization of the new host (Ribet and Cossart, 2015; Sheppard et al., 2018; Montoya et al., 2021). A comprehensive study based on codon adaptation index reported that the natural selection and host adaptation have been occurred in SARS-CoV-2 (Roy et al., 2021). Similar finding had also been reported by Lu et al., 2020. Therefore, in summary, this study shows that coronaviruses belonging to Sarbecovirus in bat could be the origin of SARS-CoV-2.

In addition to genome phylogeny, gene tree analysis was also conducted as it provides a more reliable basis for studying species evolution. Five gene trees namely Orf1ab, Spike, Envelope, Membrane, and Nucleocapsid of the *Betacoronaviruses* were reconstructed for gene tree analysis (Fig. 4 and Figs. S2-S5). Except Nucleocapsid gene tree (Fig. S5), other four gene trees have shown that the five subgenera clustered according to their cladistic relations where Embecovirus clade appeared as a basal radiation of the *Betacoronavirus* gene trees. Further, these gene trees were in concordance with the genome tree. The topological difference of Nucleocapsid gene tree with the *Betacoronavirus* genome/species tree might be possible as gene tree differs from species tree for various analytical and/or biological reasons (Degnan et al., 2009; Som, 2013; 2015). Further, analysis on the gene trees found, except Envelope gene tree, other four gene trees exhibited bat-CoV-RaTG13 is the closest relative of SARS-CoV-2 followed by pangolin-CoV as found in the genome tree analysis (Figs. 4, S2, S3, S5). Different evolutionary pattern of Envelope gene tree is probably due to stochastic error as its length is very small (average length 75 amino acids) (Som, 2015). Further analysis of the gene trees found though subgenera-wise four gene trees are similar, but within subgenera there are widespread phylogenetic inconguences (Jeffroy et al., 2006). This result led us to hypothesize that recombination events had occurred among *Betacoronaviruses* in the past that are caused to evolve new strains including the emergence of pathogenic lineage like SARS-CoV-2.

**Figure 4:**
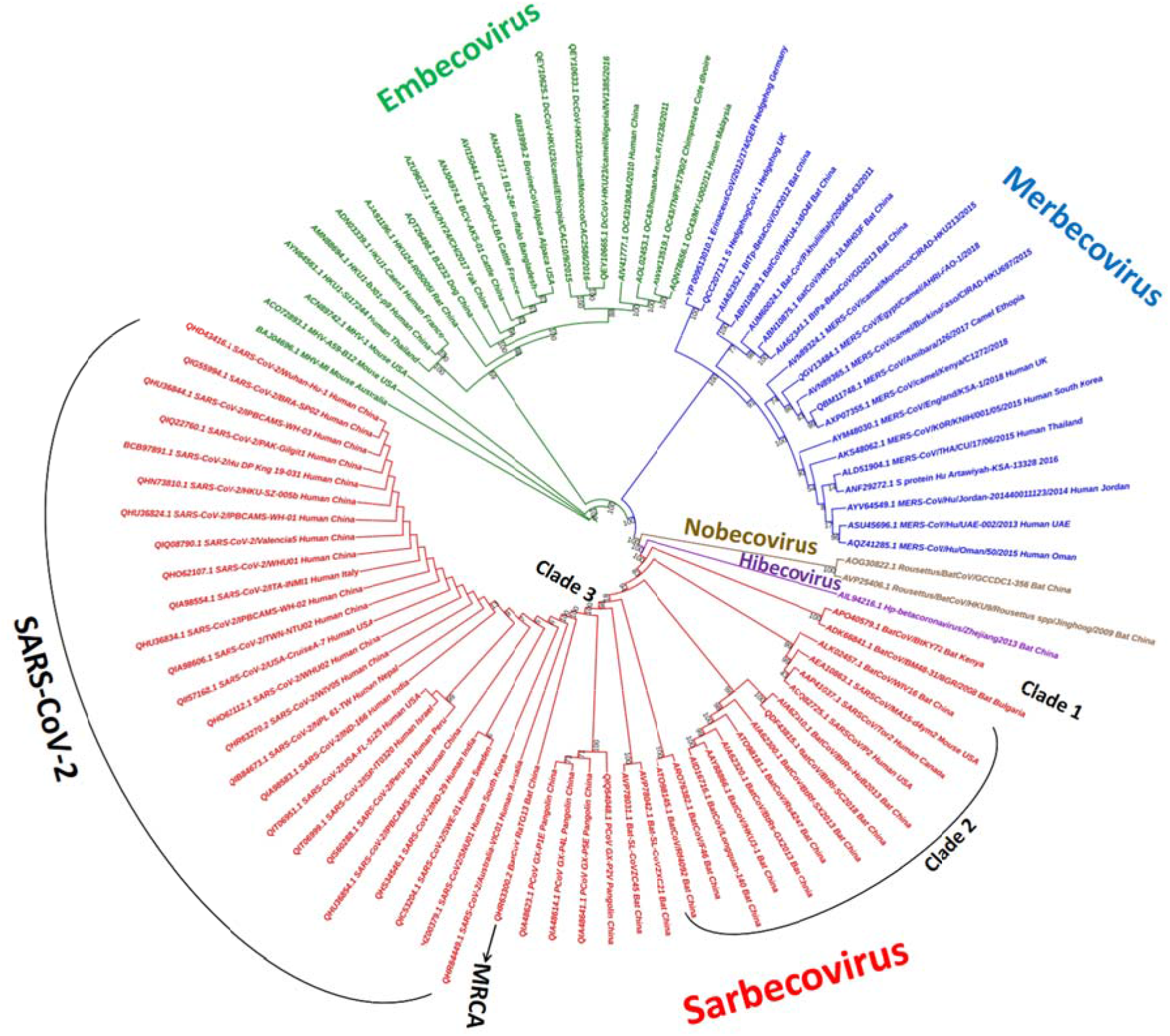
Spike (S) gene phylogeny. Alignment consists of 1,621 aligned amino acid characters (1,071bp are completely aligned characters). Tree was reconstructed using ML method and WAG+I+G4 model of protein evolution along with 1000 bootstrap replicates. Three distinct clades of sarbecovirus and most recent common ancestor (MRCA) of SARS-CoV-2 are depicted.

Accordingly, we conducted both genome and gene recombination analysis of the *Betacoronaviruses* using RDP5 package (Martin et al., 2015). The genome recombination analysis detected 21 putative recombination signals (Table 1).

**Table1:**
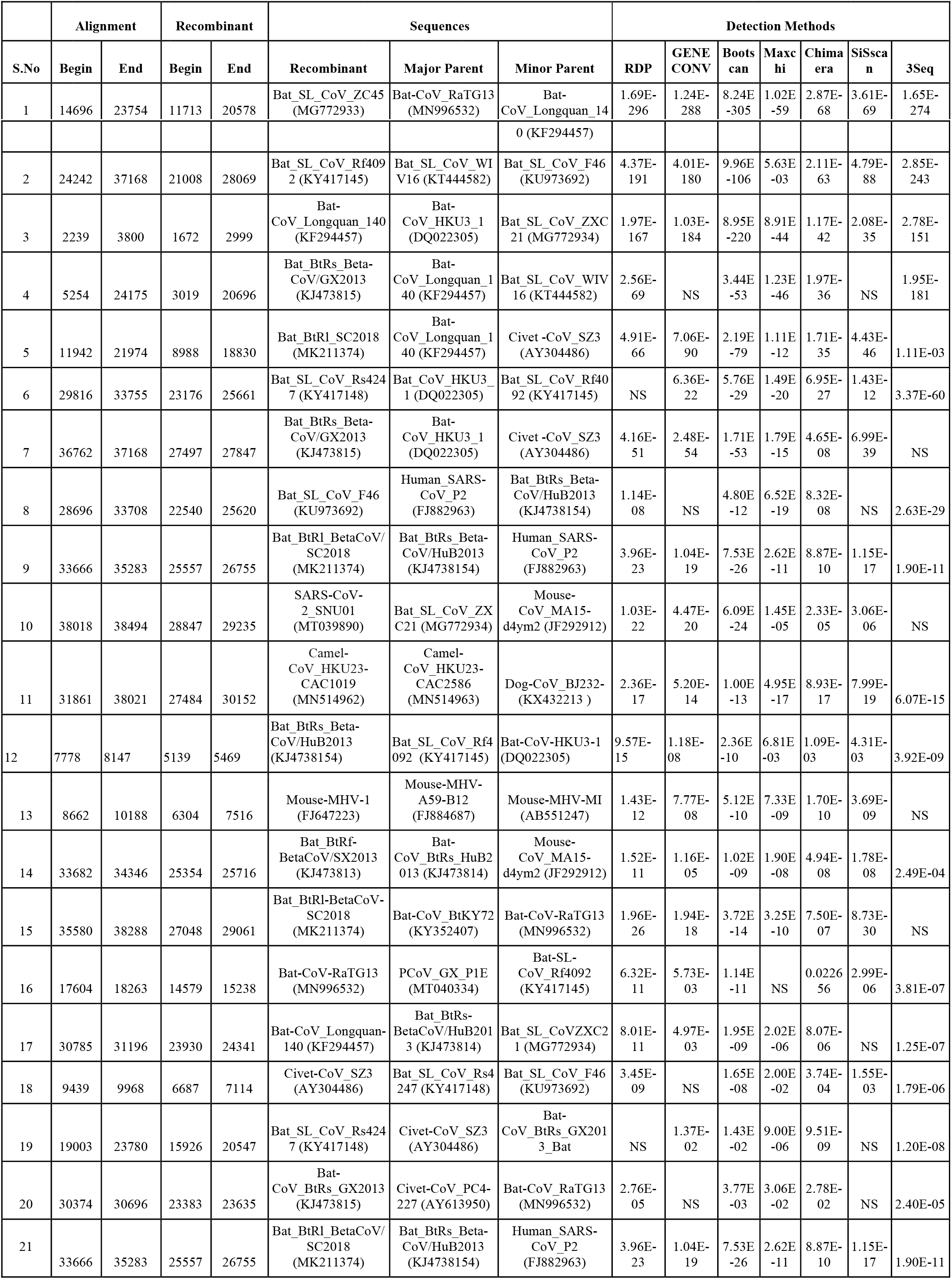
Detected recombination events in the *Betacoronavirus* genomes with position of break and endpoints, and major and minor parents. Details of genome recombination analysis are given in the text.

| S.No | Alignment |  | Recombinant |  | Sequences |  |  | Detection Methods |  |  |  |  |  |  |
| --- | --- | --- | --- | --- | --- | --- | --- | --- | --- | --- | --- | --- | --- | --- |
|  | Begin | End | Begin | End | Recombinant | Major Parent | Minor Parent | RDP | GENE CONV | Boots can | Maxc hi | Chima era | SiSca n | 3Seq |
| 1 | 14696 | 23754 | 11713 | 20578 | Bat_SL_CoV_ZC45 (MG772933) | Bat-CoV_RaTG13 (MN996532) | Bat-CoV_Longquan_14 | 1.69E-296 | 1.24E-288 | 8.24E-305 | 1.02E-59 | 2.87E-68 | 3.61E-69 | 1.65E-274 |
|  |  |  |  |  |  |  | 0 (KF294457) |  |  |  |  |  |  |  |
| 2 | 24242 | 37168 | 21008 | 28069 | Bat_SL_CoV_Rf409<br>2 (KY417145) | Bat_SL_CoV_WI<br>V16 (KT444582) | Bat_SL_CoV_F46<br>(KU973692) | 4.37E-191 | 4.01E-180 | 9.96E-106 | 5.63E-03 | 2.11E-63 | 4.79E-88 | 2.85E-243 |
| 3 | 2239 | 3800 | 1672 | 2999 | Bat-CoV_Longquan_140<br>(KF294457) | Bat-CoV_HKU3_1<br>(DQ022305) | Bat_SL_CoV_ZXC<br>21 (MG772934) | 1.97E-167 | 1.03E-184 | 8.95E-220 | 8.91E-44 | 1.17E-42 | 2.08E-35 | 2.78E-151 |
| 4 | 5254 | 24175 | 3019 | 20696 | Bat_BtRs_Beta-CoV/GX2013<br>(KJ473815) | Bat-CoV_Longquan_1<br>40 (KF294457) | Bat_SL_CoV_WIV<br>16 (KT444582) | 2.56E-69 | NS | 3.44E-53 | 1.23E-46 | 1.97E-36 | NS | 1.95E-181 |
| 5 | 11942 | 21974 | 8988 | 18830 | Bat_BtR1_SC2018<br>(MK211374) | Bat-CoV_Longquan_1<br>40 (KF294457) | Civet -CoV_SZ3<br>(AY304486) | 4.91E-66 | 7.06E-90 | 2.19E-79 | 1.11E-12 | 1.71E-35 | 4.43E-46 | 1.11E-03 |
| 6 | 29816 | 33755 | 23176 | 25661 | Bat_SL_CoV_Rs424<br>7 (KY417148) | Bat_CoV_HKU3_1<br>(DQ022305) | Bat_SL_CoV_Rf40<br>92 (KY417145) | NS | 6.36E-22 | 5.76E-29 | 1.49E-20 | 6.95E-27 | 1.43E-12 | 3.37E-60 |
| 7 | 36762 | 37168 | 27497 | 27847 | Bat_BtRs_Beta-CoV/GX2013<br>(KJ473815) | Bat-CoV_HKU3_1<br>(DQ022305) | Civet -CoV_SZ3<br>(AY304486) | 4.16E-51 | 2.48E-54 | 1.71E-53 | 1.79E-15 | 4.65E-08 | 6.99E-39 | NS |
| 8 | 28696 | 33708 | 22540 | 25620 | Bat_SL_CoV_F46<br>(KU973692) | Human_SARS-CoV_P2<br>(FJ882963) | Bat_BtRs_Beta-CoV/HuB2013<br>(KJ4738154) | 1.14E-08 | NS | 4.80E-12 | 6.52E-19 | 8.32E-08 | NS | 2.63E-29 |
| 9 | 33666 | 35283 | 25557 | 26755 | Bat_BtR1_BetaCoV/<br>SC2018<br>(MK211374) | Bat_BtRs_Beta-CoV/HuB2013<br>(KJ4738154) | Human_SARS-CoV_P2<br>(FJ882963) | 3.96E-23 | 1.04E-19 | 7.53E-26 | 2.62E-11 | 8.87E-10 | 1.15E-17 | 1.90E-11 |
| 10 | 38018 | 38494 | 28847 | 29235 | SARS-CoV-<br>2_SNU01<br>(MT039890) | Bat_SL_CoV_ZX<br>C21 (MG772934) | Mouse-CoV_MA15-d4ym2 (JF292912) | 1.03E-22 | 4.47E-20 | 6.09E-24 | 1.45E-05 | 2.33E-05 | 3.06E-06 | NS |
| 11 | 31861 | 38021 | 27484 | 30152 | Camel-CoV_HKU23-CAC1019<br>(MN514962) | Camel-CoV_HKU23-CAC2586<br>(MN514963) | Dog-CoV_BJ232-<br>(KX432213 ) | 2.36E-17 | 5.20E-14 | 1.00E-13 | 4.95E-17 | 8.93E-17 | 7.99E-19 | 6.07E-15 |
| 12 | 7778 | 8147 | 5139 | 5469 | Bat_BtRs_Beta-CoV/HuB2013<br>(KJ4738154) | Bat_SL_CoV_Rf4<br>092 (KY417145) | Bat-CoV-HKU3-1<br>(DQ022305) | 9.57E-15 | 1.18E-08 | 2.36E-10 | 6.81E-03 | 1.09E-03 | 4.31E-03 | 3.92E-09 |
| 13 | 8662 | 10188 | 6304 | 7516 | Mouse-MHV-1<br>(FJ647223) | Mouse-MHV-A59-B12<br>(FJ884687) | Mouse-MHV-MI<br>(AB551247) | 1.43E-12 | 7.77E-08 | 5.12E-10 | 7.33E-09 | 1.70E-10 | 3.69E-09 | NS |
| 14 | 33682 | 34346 | 25354 | 25716 | Bat_BtRf-BetaCoV/SX2013<br>(KJ473813) | Bat-CoV_BtRs_HuB2<br>013 (KJ473814) | Mouse-CoV_MA15-d4ym2 (JF292912) | 1.52E-11 | 1.16E-05 | 1.02E-09 | 1.90E-08 | 4.94E-08 | 1.78E-08 | 2.49E-04 |
| 15 | 35580 | 38288 | 27048 | 29061 | Bat_BtR1-BetaCoV-SC2018<br>(MK211374) | Bat-CoV_BtKY72<br>(KY352407) | Bat-CoV-RaTG13<br>(MN996532) | 1.96E-26 | 1.94E-18 | 3.72E-14 | 3.25E-10 | 7.50E-07 | 8.73E-30 | NS |
| 16 | 17604 | 18263 | 14579 | 15238 | Bat-CoV-RaTG13<br>(MN996532) | PCoV_GX_P1E<br>(MT040334) | Bat-SL-CoV_Rf4092<br>(KY417145) | 6.32E-11 | 5.73E-03 | 1.14E-11 | NS | 0.0226<br>56 | 2.99E-06 | 3.81E-07 |
| 17 | 30785 | 31196 | 23930 | 24341 | Bat-CoV_Longquan-140 (KF294457) | Bat_BtRs-BetaCoV/HuB201<br>3 (KJ473814) | Bat_SL_CoVZXC2<br>1 (MG772934) | 8.01E-11 | 4.97E-03 | 1.95E-09 | 2.02E-06 | 8.07E-06 | NS | 1.25E-07 |
| 18 | 9439 | 9968 | 6687 | 7114 | Civet-CoV_SZ3<br>(AY304486) | Bat_SL_CoV_Rs4<br>247 (KY417148) | Bat_SL_CoV_F46<br>(KU973692) | 3.45E-09 | NS | 1.65E-08 | 2.00E-02 | 3.74E-04 | 1.55E-03 | 1.79E-06 |
| 19 | 19003 | 23780 | 15926 | 20547 | Bat_SL_CoV_Rs424<br>7 (KY417148) | Civet-CoV_SZ3<br>(AY304486) | Bat-CoV_BtRs_GX201<br>3_Bat | NS | 1.37E-02 | 1.43E-02 | 9.00E-06 | 9.51E-09 | NS | 1.20E-08 |
| 20 | 30374 | 30696 | 23383 | 23635 | Bat-CoV_BtRs_GX2013<br>(KJ473815) | Civet-CoV_PC4-227 (AY613950) | Bat-CoV_RaTG13<br>(MN996532) | 2.76E-05 | NS | 3.77E-03 | 3.06E-02 | 2.78E-02 | NS | 2.40E-05 |
| 21 | 33666 | 35283 | 25557 | 26755 | Bat_BtR1_BetaCoV/<br>SC2018<br>(MK211374) | Bat_BtRs_Beta-CoV/HuB2013<br>(KJ4738154) | Human_SARS-CoV_P2<br>(FJ882963) | 3.96E-23 | 1.04E-19 | 7.53E-26 | 2.62E-11 | 8.87E-10 | 1.15E-17 | 1.90E-11 |

A recombination event was reported when five out of seven methods detected it. Recombination results show that major recombination events took place between bat coronaviruses belonging to the subgenus Sarbecoviruses. A recent study by Boni et al (2020) also reported the Serbicoviruses lineage undergoes frequent recombination. For further insights, we compared SARS-CoV-2 Hong Kong (HKU_SZ_005b) genome sequence with four closely related SARS-CoVs namely Bat-CoV-RaGT13, Bat-SL-CoVZC45, Bat-SL-CoVZXC21, and Pangolin-CoV-GX-P5E using simplot analysis (Fig. 5). Simplot exhibits that bat-CoV-RaTG13 show the highest similarity with SARS-CoV-2 genome including exchange of genetic materials at the different regions as shown in Figure 5. We classified the whole genomes into four regions (Regions1-4).

**Figure 5:**
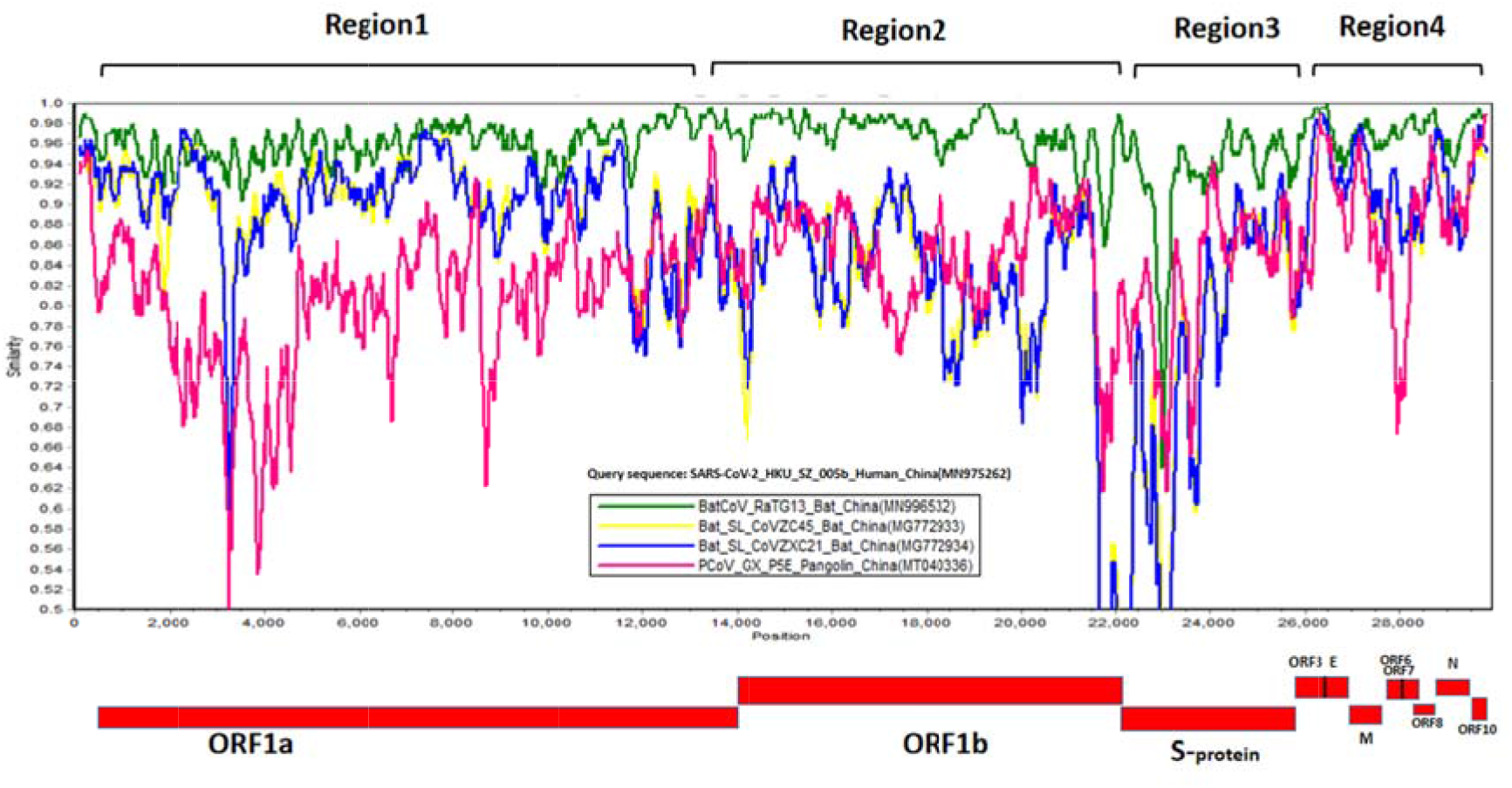
Similarity plot (Simplot) of SARS-CoV-2 HKU-China and its comparison with other Coronaviruses (Green, Bat-CoV-RaGT13; Pink, Pangolin-CoV-GX-P5E; Yellow Bat-SL-CoVZC45: and Blue, Bat-SL-CoVZXC21). Simplot depicts the Bat and Pangolin CoVs showing recombination. Four different regions (regions 1-4) from the genomes showing recombination were highlighted.

In region 1 (which mostly covers ORF1a gene), we observed highest genetic divergence between pangolin and SARS-CoV-2 strains, and bat to bat recombinationevents were frequent. In region2 (ORF1b gene), recombination events mostly took place between bat and pangolin strains. In region3 (Spike gene), bat-CoV-RaTG13 genome shows divergence with SARS-CoV-2 genome and there is a good number of genetic recombination among the bat and pangolin strains. In region4 (E, M, N and ORF3/6-8/10 genes), all strains show high similarity and a few number of recombination events with the SARS-CoV-2 strain. Further, gene recombination analysis found that there are highest recombination events in spike protein (spotted nine events) followed by Orf1ab protein (six events). Membrane and Nucleocaspid proteins reported few recombination events and envelope protein did not show any recombination event. Overall, recombination results support our phylogenetic inference and suggest that the origin of SARS-CoV-2 is the results of ancestral intra-species recombination events between bat SARS-CoVs (Flores-Alanis et al., 2020; Li et al., 2020). Details of recombination analysis are given in Table 1.

Further we measured the genetic variation of bat-CoV-RaTG13 and pangolin-CoV-GX-P5E sequences with respect to SARS-CoV-2 Wuhan-Hu-1 strain, and found that spike protein has highest genetic variation 3% and 7 % respectively (Table 2).

**Table 2:** Homology and genetic variations in different genomic regions of SARS-CoV-2 Wuhan (MN908947) with respect to Bat-CoV-RaTG13 (MN996532) and Pangolin-CoV-GX-P5E (MT040336).

| Strain | Envelop protein |  | Membrane protein |  | Spike protein |  | Nucleocapsid protein |  |
| --- | --- | --- | --- | --- | --- | --- | --- | --- |
|  | Homology | Genetic variation | Homology | Genetic variation | Homology | Genetic variation | Homology | Genetic variation |
| Bat_Ra TG13 | 100% | 0% | 98% | 02% | 97% | 03% | 99% | 01% |
| PCoV_GX-P5E | 100% | 0% | 98% | 02% | 92% | 08% | 93% | 07% |

Major genetic variations in spike protein seemed essential for the transition from animal-to-human transmission to human-to-human transmission of SARS-CoV-2 (Su et al., 2016; Luk et al. 2019; Jaimes et al., 2020; Mondal et al., 2021). We further did MSA of the spike protein sequences and observed that the insertion of the novel amino acids “PRRA” in the spike protein of SARS-CoV-2 (Fig. 6). A number of studies also reported/observed the insertion of “PRRA” residues in the spike protein of SARS-COV-2 (Budhraja et al., 2021; Coutard et al., 2020; Wang et al., 2020; Zhang et al., 2021).

**Figure 6:**
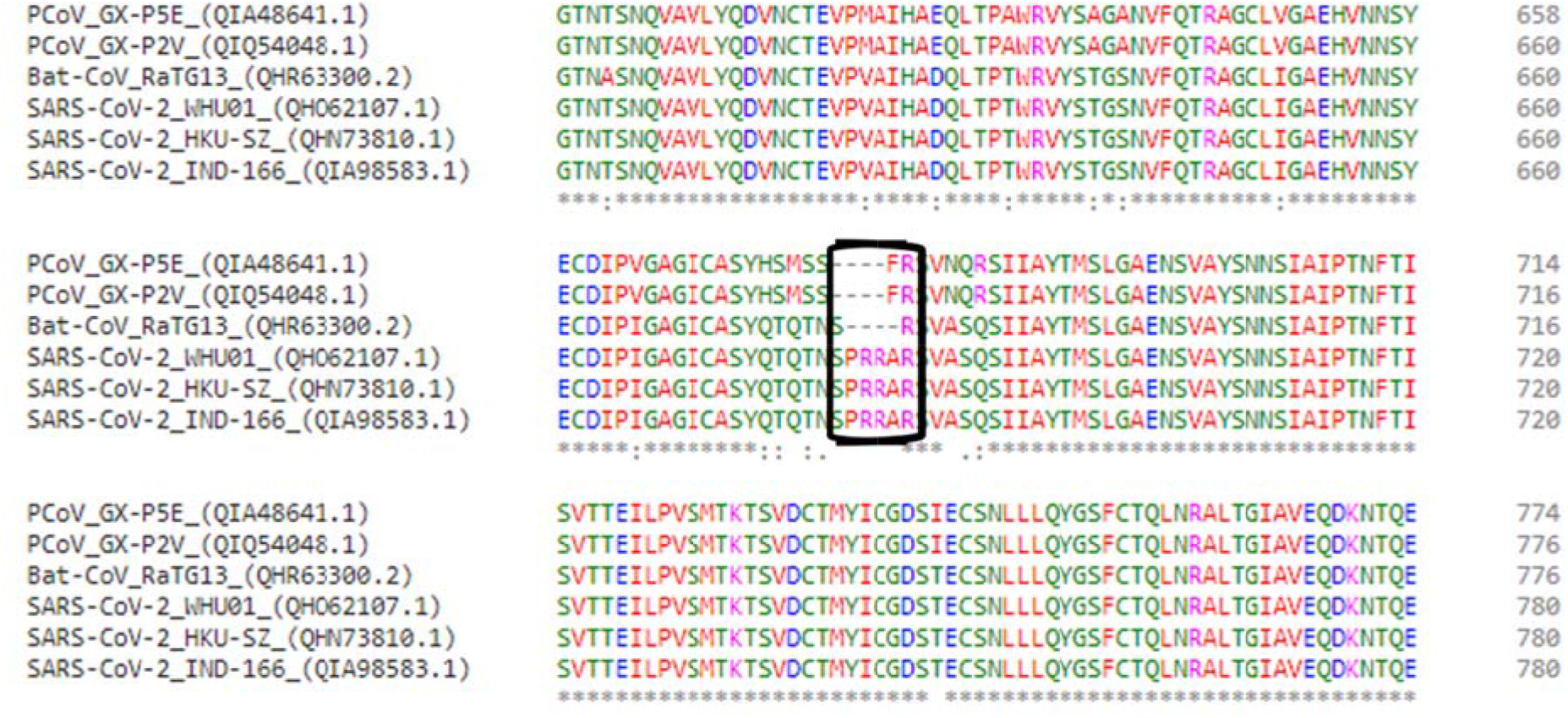
Multiple sequence alignment of spike (S) protein consisting of six strains (three SARS-CoV-2s and three closest CoV strains from bat and pangolin).

The “PRRA” insertion at the S1/S2 junction site which induces a furin cleavagemotif needs to be investigated. Therefore, further detailed study on these residues would be required to shed light on molecular mechanism of interaction between SARS-CoV-2 and host cells.

On the basis of MSA result, we compared the structure of spike protein of SARS-CoV-2 (PDB: 6XLU) with bat-CoV-RaTG13 (PDB: 6ZGF) and pangolin-CoV-GX-P5E (modeled protein) (Figure 7). The spike protein is a complex trimeric protein and monomer was used for structure comparison. It has two main units S1 and S2. The S1 subunit recognizes and binds to the host receptor enzyme via receptor-binding domains (RBDs) while the S2 subunit helps in fusion of viral cell membrane to host cell (Jaimes et al., 2020; Rehman et al., 2020; Mitra et al. 2020). We found that structurally the spike protein of pangolin-CoV-GX-P5E is more diverse compared to SARS-CoV-2 (rmsd value 2.766 Å) while the bat-CoV-RaTG13 spike protein shows similarity to SARS-CoV-2 with rmsd 2.059 Å (Figure 7).

**Figure 7:**
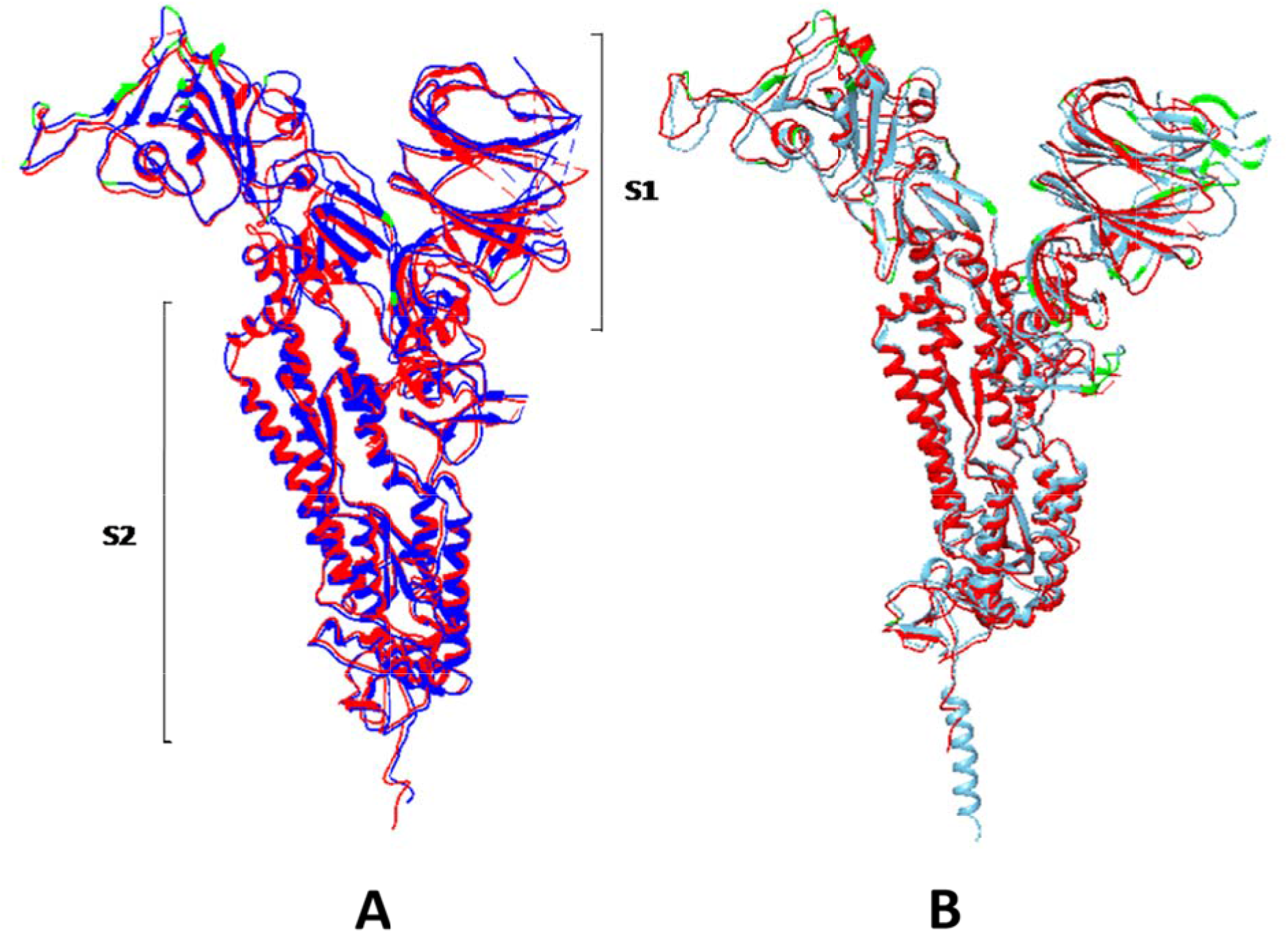
Structural representation of spike glycoprotein (S) and their comparison. Spike protein monomer superimposed structure of **(A)** SARS-CoV-2 Wuhan-Hu-1 (red) and bat-CoV-RaTG13 (blue), **(B)** SARS-CoV-2 Wuhan-Hu-1 & Pangolin-CoV-GX-P5E (sky blue). The green coloured highlighted regions represent the mutated amino acid residues.

It was observed that the bat-CoV-RaTG13 shows high number of mutations at one of the RBD (spotted 27 mutations: shown in green colour in Figure 7) while the pangolin-CoV-GX-P5E shows mutations at both the RBDs of S1 subunit (a total of 85 mutations). The changes in spike proteins have impact on the interaction of pathogen and host (Li 2015; Huang et al., 2020). Thus these mutations were probably responsible for the adaptation of SARS-CoV-2 into human systems. A number of studies reported that the mutations in spike protein of SARS-CoV-2 facilitate its adaptation into humans (Choe et al., 2021; Isabel et al., 2021; Zhang et al., 2021). The insertion of the four amino acids “PRRA” found in the MSA represents an extended loop between the two paralllel β-sheets (S1/S2 cleavage site). This cleavage point between the receptor binding domain (S1) and fusion peptide (S2) mediate cell-cell fusion and entry into human cell (Andersen et al. 2020; Mitra et al. 2020). Thus structural analysis supports MSA results and suggests that SARS-Cov-2 is adapted to infect human systems.

## 4. Concluding remarks

Outbreak of SARS-CoV-2 is the third documented spillover of an animal coronavirus to humans in only two decades that has resulted in a major pandemic. In quest of the origin, evolution and adaptation of SARS-CoV-2, our analysis suggested that the probable origin of SARS-CoV-2 is the results of ancestral intra-species recombination events between bat coronaviruses belonging to Sarbecovirus subgenus and the insertion of the four amino acids “PRRA” in the spike protein of SARS-CoV-2 along with high number of mutations at one of its receptor-binding domain are probably responsible for the adaptation of SARS-CoV-2 into humans systems. Thus, our findings add strength to the existing knowledge on the origin and adaptation of SARS-CoV-2. Further a detailed mechanistic understanding of molecular mechanisms of interaction between SARS-CoV-2 and host cells is crucial for more effective vaccine design and predicting future pandemics.

## Conflicts of interest

The authors declare that they have no conflict of interest.

## Ethical approval

This article does not contain any studies with human participants or animals performed by any of the authors.

## Acknowledgements

Thanks to Arindam Ghosh for useful discussions. This work was partly supported by the Department of Biotechnology (DBT) and University Grants Commission (UGC), India.

## Supplementary data

**Supplementary File S1:** Details of the 162 *Orthocoronavirinae* genomes and four outgroup sequences used in this study.

**Figure S2: Orf1ab gene tree.** Alignment consists of 8,152bp aligned amino acid characters (6,276bp are completely aligned characters). Tree was reconstructed using ML method and LG+I+G4 model of protein evolution along with 1000 bootstrap replicates.

**Figure S3: Membrane (M) gene tree.** Alignment consists of 233bp aligned amino acid characters (213bp are completely aligned characters). Tree was reconstructed using ML method by and LG+G4 model of protein evolution along with 1000 bootstrap replicates.

**Figure S4: Envelope (E) gene tree.** Alignment consists of 90bp aligned amino acid characters (74bp are completely aligned characters). Tree was reconstructed using ML method and JTT+I+G4 model of protein evolution along with 1000 bootstrap replicates.

**Figure S5: Nucleocaspid (N) gene tree.** Alignment consists of 547bp aligned amino acid characters (343 are completely aligned characters). Tree was reconstructed using ML and LG+I+G4 model of protein evolution along with 1000 bootstrap replicates.

## Notes

### Competing Interest Statement

The authors have declared no competing interest.

